# The evolution of the human DNA replication timing program

**DOI:** 10.1101/2022.08.09.503365

**Authors:** Alexa N. Bracci, Anissa Dallmann, Qiliang Ding, Melissa J. Hubisz, Madison Caballero, Amnon Koren

**Affiliations:** Department of Molecular Biology and Genetics, Cornell University, Ithaca, NY 14853, USA; Bioinformatics Facility, Institute of Biotechnology, Cornell University, Ithaca, NY 14853, USA

## Abstract

DNA is replicated according to a defined spatiotemporal program that is linked to both gene regulation and genome stability. The evolutionary forces that have shaped replication timing programs in eukaryotic species are largely unknown. Here, we studied the molecular causes and consequences of replication timing evolution across 94 humans, 95 chimpanzees, and 23 rhesus macaques. Replication timing differences recapitulated the species’ phylogenetic tree, suggesting continuous evolution of the DNA replication timing program in primates. Hundreds of genomic regions had significant replication timing variation between humans and chimpanzees, of which 66 showed advances in replication origin firing in humans while 57 were delayed. Genes overlapping these regions displayed correlated changes in expression levels and chromatin structure. Many human-chimpanzee variants also exhibited inter-individual replication timing variation, pointing to ongoing evolution of replication timing at these loci. Association of replication timing variation with genetic variation revealed that DNA sequence evolution can explain replication timing variation between species. Taken together, DNA replication timing shows substantial and ongoing evolution in the human lineage that is driven by sequence alterations and impacts regulatory evolution.

## Introduction

Understanding of human specific phenotypes and their evolution has primarily focused on the comparison of individual genes or sequence elements, and their regulation, between humans and closely related species [1]. Humans and chimpanzees are approximately 99% identical at the single nucleotide level, yet have undergone extensive phenotypic divergence [2]. This has increasingly been attributed to regulatory evolution, including gene expression, which has been associated with brain, skeletal, and other phenotypes [3–6]. An understudied form of genome regulation, with potential impact on regulatory and sequence evolution, is the spatiotemporal program of DNA replication.

Genome replication is accomplished by replication origins that fire at different times during S phase, resulting in a defined pattern of DNA replication timing. Early DNA replication is associated with high gene density, open chromatin, and active transcription [7], while later replicating regions typically exhibit higher frequencies of single nucleotide mutations and polymorphisms [8–10]. Replication timing thus bridges between genome regulation and evolution. As a corollary, understanding the evolution of replication timing can reveal the selective forces that have shaped particular replication programs, inform mechanisms of replication timing regulation, and uncover impacts of replication timing on sequence, molecular and phenotypic evolution.

Only a handful of studies have compared replication timing across species. Studies in yeast suggested that replication origins dynamically gain and lose activity during evolution [11] and that conserved early replication, in particular of histone genes, is required for high gene expression levels [12]. In contrast, replication timing has been shown to be highly conserved between corresponding cell types of humans and mice despite extensive genome rearrangements [13, 14], while a more recent study suggested the presence of both conserved and species-specific replication timing regions among five primate species [15]. Importantly, previous studies have been under-powered to identify the genetic changes that drive replication timing evolution nor its potential impacts on regulatory and sequence evolution.

Here, we address the causes and consequences of replication timing evolution by profiling a large number of humans, chimpanzees, and rhesus macaques. We find that replication timing has continuously evolved across these species at hundreds of locations. Comparison to intra-species variation and sequence polymorphisms within species and divergence between species revealed the genetic basis of a subset of replication origins that have gained or lost activity during evolution. On the other hand, analysis of gene expression and chromatin structure suggests a complex relationship between the evolution of replication timing and gene regulation. Overall, this study advances our knowledge of how replication timing evolves, the association of replication timing with genome regulation and transcription, and the determinants of replication timing evolution.

## Results

### High resolution DNA replication timing profiles across humans, chimpanzees, and rhesus macaques

To study the evolution of DNA replication timing across primates, we sequenced the genomes of 90 chimpanzee lymphoblastoid cell lines (LCL), 23 rhesus macaque LCLs, and seven chimpanzee induced pluripotent stem cell lines (iPSC), along with 88 human LCLs and eight human iPSCs. We aligned each species’ sequencing reads to its own reference genome and inferred DNA replication timing from read depth fluctuations across chromosomes [16–18] (see Methods). Our method of inferring replication timing from whole genome sequence data was particularly suited to this task, as chimpanzee material is scarcely available for the experimental manipulations required by other approaches (e.g. Repli-seq; [19]). One chimpanzee LCL, one chimpanzee iPSC and two human iPSC were filtered due to low data quality. Read depth fluctuations showed long-range continuity along chromosomes (autocorrelation) consistent with DNA replication, and LCL data resolution was further improved using principal component (PC)-regression [18] (Figure S1 A-E). The resulting replication timing profiles were highly consistent across samples within each species (human LCLs r = 0.94—0.99; chimpanzee LCLs r = 0.84–1; rhesus LCLs r = 0.97–1; human iPSCs r = 0.91–0.97; chimpanzee iPSCs r = 0.96–0.97) (Figure 1 A-E; S2 A-D).

**Figure 1.**
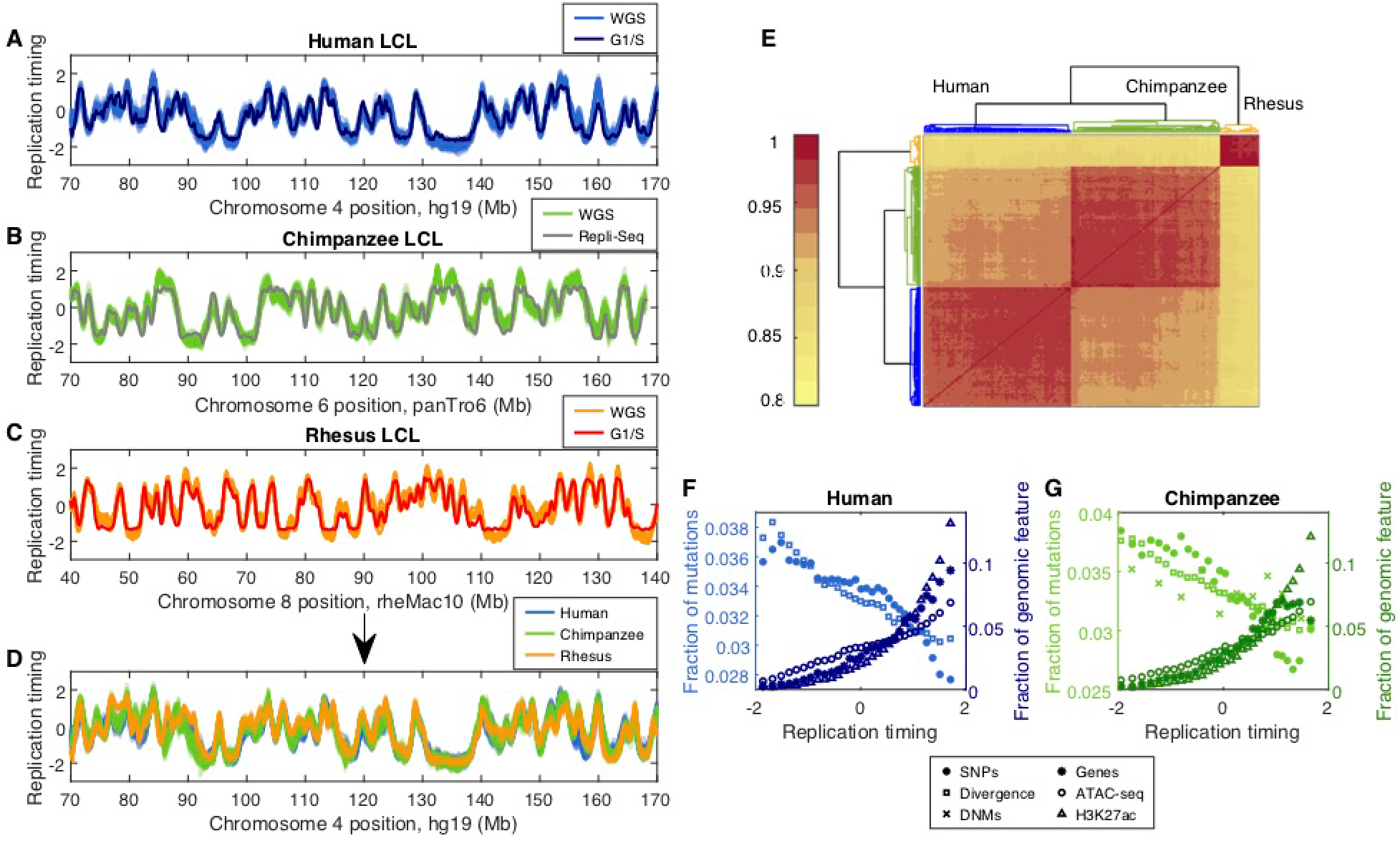
Replication timing evolution in primate species. (A-C) Replication timing was inferred from read depth fluctuations in whole genome sequencing (WGS) data of human, chimpanzee, and rhesus macaque LCLs. Units are standard deviation from an autosome-wide mean of 0. Shown for comparison are a consensus G1/S profile for human LCLs (A) [9], a G1/S rhesus macaque LCL profile (generated in this study; C), and a chimpanzee LCL replication profile generated using Repli-Seq (B) [15]. (D) Human, chimpanzee, and rhesus macaque replication timing profiles, plotted on human genomic coordinates (hg19; see also Figure S3), show conservation of the replication timing program. (E) Hierarchical clustering of human, chimpanzee, and rhesus macaque LCL Pearson correlation values. Replication timing is highly consistent within species and, while is largely conserved among species, exhibits significant inter-species variation that corresponds to the evolutionary divergence of primates. (F-G) SNPs, human-chimpanzee divergent sites, and de novo mutations (DNMs) are enriched at late-replicating DNA, while protein coding genes and marks of accessible chromatin (ATAC-seq peaks and H3K27ac ChIP-seq peaks) are enriched at early-replicating DNA. Fraction of human (F) or chimpanzee (G) genomic features in 30 replication timing bins per species. DNM rate was calculated in 10 replication timing bins (fraction is 3x).

We further validated the replication timing profiles in three ways. First, we measured replication timing by sorting and sequencing G1 and S phase cells of select samples [9], which provided replication timing profiles highly correlated to those generated without cell sorting (human LCL mean r = 0.97; chimpanzee iPSC mean r = 0.87; rhesus LCL mean r = 0.95) (Figure 1A, 1C; S2B). Second, we compared the chimpanzee LCL samples to a previously published chimpanzee replication timing profile generated using Repli-seq [15] (mean r = 0.90) (Figure 1B). Finally, as we showed previously in humans [20], replication of the X chromosomes was delayed and less structured in chimpanzee LCL females compared to males (Figure S1 G-J). Together, these results demonstrate that the replication timing profiles of all three species are of high quality and reproducibility.

Next, we compared genome-wide replication timing to human-chimpanzee divergence, single nucleotide polymorphisms (human dbSNP 153 common, n=9,585,612; chimpanzee dbSNP, n=1,468,866), and somatic cell line mutations (identified as de novo mutations in chimpanzee trios; see Methods) and found that all were enriched at late-replicating genomic regions in both humans and chimpanzees (Figure 1 F, G). On the other hand, gene density, gene expression levels [21–24], ATAC-seq [25], and H3K27ac ChIP-seq [23] were all enriched at, or correlated with, early-replicating genomic regions in both humans and chimpanzees (Figure 1 F, G; S2 J). These genome-wide trends were also replicated in human and chimpanzee iPSCs [21, 22] (Figure S2 H-J), overall supporting the conservation of genomic features associated with replication timing across cell types and species.

### Substantial variation in replication timing between species

To compare replication timing profiles between species, we converted the chimpanzee and rhesus macaque replication timing data to human genome coordinates (see Methods); these conversions had a minimal effect on the structures of the replication profiles (Figure S3 A, B). We found that replication timing was highly conserved across species (Figure 1 D, E; S2 C, D) and that replication timing variation was greatest between cell types (LCL and iPSC; mean r = 0.69) (Figure S2 E-G). Nonetheless, there were also clear inter-species differences within the same cell type. Hierarchical clustering of replication timing values across samples recapitulated the phylogenetic tree for these three species (Figure 1E; S2D), suggesting that replication timing has evolved continuously across the primate lineage, primarily in a cell-type-specific manner.

We systematically searched for specific differences in replication timing between humans and chimpanzees, separately for LCLs and iPSCs, using sliding ANOVA tests with a Bonferroni corrected p-value threshold of 8.7×10^-7^ (see Methods). For the X chromosome, males and females were considered separately. We identified 858 autosomal regions where human and chimpanzee LCLs significantly differed in replication timing. These regions covered 1.1 Mb on average and cumulatively spanned 980 Mb (36.6% of the analyzable autosomes). Similarly, we identified 47 variant regions on the X chromosome in females and 39 in males (1.4 and 1.2 Mb on average, spanning a total of 64 (42.9% of the analyzable X chromosome) and 45 Mb (29.9%) in females and males, respectively). In iPSCs, we identified 704 autosomal variant regions covering 1.1 Mb on average and cumulatively spanning 797 Mb (29.8% of the autosomes), likely less than in LCLs due to the more limited sample size.

A majority of the human-chimpanzee variant regions occurred at peaks in the replication timing profiles (LCL: 620/944, 65.7%; iPSC: 476/704, 67.6%), suggesting that a major source of replication timing variation is changes in replication origin (or origin cluster) activity. Extending from this observation, and under the assumption that changes in origin activity are the most likely explanation for the large replication timing variants that we observed, we designated the center of the peak as the most likely source of replication timing variation within each region (see Methods). This was only applied to variant regions that contained replication timing peaks, while 134 LCL variants with more than one peak were separated into several independent variant regions. Overall, we called 731 LCL and 557 iPSC replication timing variants that each contained one putative source site (Figure 2; S4; S5; Table S1), and utilized them for downstream analyses. These variants covered on average 1.2 Mb in LCLs and 1.1 Mb in iPSCs, and had an average magnitude of replication timing difference between humans and chimpanzees of 0.4 standard deviations in LCLs (Figure S4 B, C) and 0.5 in iPSCs. The majority of these variant regions were earlier replicating in humans compared to chimpanzees, in both LCLs (57.0%; Binomial test p=1.2×10^-4^; Figure 2E; S4B) and iPSCs (53.9%; p=0.08). Consistent with these variants containing replication profile peaks, the distribution of replication timing at variants was skewed towards early replication in both humans and chimpanzees (Figure S4A). The fraction of replication timing peaks that varied between humans and chimpanzees – 32.5% – was comparable to the fraction of the genome with replication timing variation; thus, the widespread evolution of replication timing is not an inflated estimate due to the broad effect of individual replication origins.

**Figure 2.**
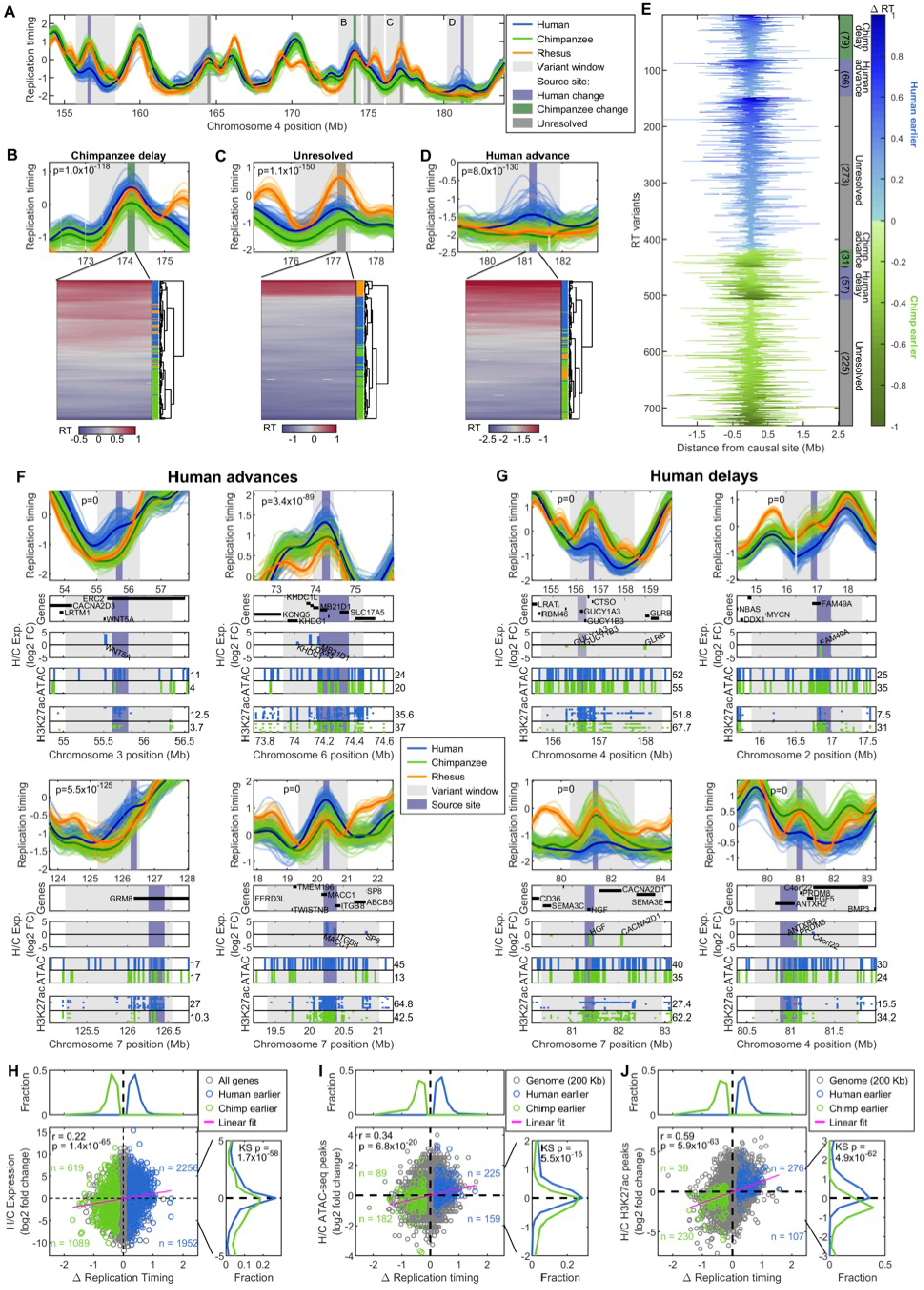
Replication timing evolution and its co-variation with gene expression and chromatin accessibility. (A) Replication timing profiles for a region of chromosome 4 for humans (n=88, blue), chimpanzees (n=89, green), and rhesus macaques (n=23, orange) along with identified human-chimpanzee replication timing variant regions (light gray) and their called source sites. Source site color indicates the lineage in which replication timing was inferred to have evolved. (B-D) Three example regions indicated in (A) are shown at greater resolution. Replication timing of each sample within the source site shown as heat maps; dendrograms: hierarchical clustering of sample replication timing similarity. The clustering demonstrates the separation of the majority (or all) of human from chimpanzee samples as well as the clustering of one (or none) of them to rhesus macaque replication timing. P-values: significance (ANOVA) of human-chimpanzee differences within the variant region. (E) Mean difference in replication timing (Δ RT) between humans and chimpanzees across each replication timing variant, centered at the source sites. Variants are sorted by being earlier in humans or chimpanzees, then by the species in which the change was inferred to have happened, and last by the magnitude of inter-species replication timing difference (Δ RT). (F-G) Examples of replication timing advances (F) and delays (G) inferred to have occurred in the human lineage. Genes, expression, ATAC-seq peaks and H3K27ac ChIP-seq data shown beneath the replication timing profiles for each variant and flanking 200 Kb. Numbers next to ATAC-seq track: the number of human and chimpanzee ATAC-seq peaks within the variant window. Numbers next to H3K27ac track: average number of human and chimpanzee H3K27ac ChIP-seq peaks within the variant window. Some gene names were removed from the Genes track for readability. (H) Differences between human and chimpanzee LCL replication timing compared to differences in gene expression, for all genes as well as genes within replication timing variants with either earlier replication timing in humans (blue) or in chimpanzees (green). Correlation coefficient (r) and p-value indicated for variant regions. Number of genes in each quadrant further demonstrates the correlation between gene expression and replication timing variation. Top and right histograms: distributions of replication timing and gene expression differences, respectively, within replication timing variant regions. (I-J) As in H, using ATAC-seq (I) or H3K27ac ChIP-seq (J) data.

In order to infer whether each replication timing change occurred in the human or chimpanzee lineage, we compared the average LCL replication timing profile for each species to the average profile of rhesus macaque as an outgroup. Specifically, we calculated the pairwise Euclidean distance between each pair of species within each variant region source site (see Methods). Human-specific replication timing changes were defined as regions in which chimpanzees and rhesus macaques were closer to each other than either were to humans, and chimpanzee-specific changes were similarly defined as cases in which humans and rhesus macaques were the most similar (see Methods). Regions with significantly different replication timing among all three species were considered unresolved for evolutionary direction. In total, we resolved 233 replication timing variant regions (out of the 731 LCL variants containing a replication timing peak), of which 123 and 110 were changes in the human and chimpanzee lineages, respectively (similar number of changes in each lineage expected based on the molecular evolutionary clock; χ^2^=0.73, df=1, p=0.39 [26]). Of these, we inferred 66 to be human advances and 57 to be human delays, while another 31 and 79 were inferred to be replication timing advances or delays, respectively, in chimpanzees (Figure 2; S4 D, E). Of the 66 human advances, 55 represented earlier activation of a shared origin while another 11 regions appeared to represent de novo evolutionary emergence of novel replication origins in the human lineage. Similarly, 55 replication origins appeared to have been delayed in their firing time in humans compared to chimpanzees, with evidence for two replication origins being entirely lost in some humans. Importantly, all human origin gains and losses were polymorphic, present in 31-72% and 7-13% of individuals, respectively (Figure S6). This suggests that these origins have been recently gained or lost and are subject to ongoing evolution in the human lineage. In comparison, we identified seven and 56 origins that have been putatively gained or lost, respectively, in virtually all human and chimpanzee samples compared to macaques (allowing up to 5% technical variation of samples). Thus, on a broader evolutionary timescale, we see compelling evidence of more substantial restructuring of the replication program in primates.

Of 731 human-chimpanzee LCL replication timing variants, 47 (6.4%) were shared in iPSCs; of these, 30 had a similar shape of replication timing profiles (by correlation; see Methods) between cell types of the same species (Figure S5 E). Thus, although variants shared across cell types have greater potential to impact species-specific phenotypic differences, most human-chimpanzee replication timing differences are cell-type-specific.

### Association of replication timing variation with gene evolution

iPSC variant regions spanned 2,801 protein coding genes (mean of 5 genes per variant), while LCL replication timing variants spanned a total of 6,156 protein coding genes (mean of 8.5 genes per variant). iPSC replication timing variant regions were enriched for genes involved in immunity and development (Table S2), some of which are known to be under adaptive evolution in humans (e.g. AKAP11, GLB1L2, SYBU, CD59, PYHIN1, PYDC2, SIGLEC9/L1, ADAM2, OVGP1, SEMG1, SEMG2, ANG) [27, 28]. Similarly, several genes inferred to be under positive selection in humans fell into LCL replication timing variant regions. These genes included several with roles in cell cycle progression (TLE6; Figure S7 H), Wnt signaling (TLE4), and sperm motility (CATSPER1, SEMG1, SEMG2), and several associated with human diseases or conditions including Usher Syndrome (USHBP1; Figure S7 I), glaucoma (RMDN2; Figure S7 J), intellectual disability (KPTN) and microcephaly (ASPM; Figure S7 K). One notable LCL variant region spanned the APOBEC cluster that includes APOBEC3A, APOBEC3B, APOBEC3C, APOBEC3D, APOBEC3F, APOBEC3G, and APOBEC3H; these genes play a role in antiviral activity and most have been under positive selection in primates [29] (Figure S7 G). APOBEC genes replicated earlier in humans, and most fell within the source site of replication timing variation.

We identified 877 protein coding genes with variable replication timing between human and chimpanzee in both LCL and iPS cells. One notable gene under positive selection in humans, PYHIN1 (IFIX), replicated later in humans compared to chimpanzees (and macaques; Figure S7 L). This gene is a known tumor suppressor, down regulation of which is associated with breast cancer [30].

As a complementary analysis, we examined replication timing evolution at various genomic elements previously described to undergo atypical rates of evolution. These included human-chimpanzee divergent sites; more specifically human accelerated regions (HARs), which are conserved in mammals yet have undergone many sequence changes in humans [31]; and regions identified as under ancient positive selection in humans (selective sweeps [32]). Conversely, we analyzed regions under evolutionary constraint: loss of function intolerant genes (gnomAD) [33]), and ultra-conserved elements (UCEs) that are completely conserved in sequence across human, mouse, and rat [34]. As expected, sites of sequence divergence (Figure 1 F-G) and HARs (although not sites of selective sweeps; Figure S7 C-D) were biased to late replication, while UCEs and loss of function intolerant genes replicated earlier than expected (compared to genes in general in the latter case; Figure S7 A-B). More significantly, we also found that divergent sites and regions under ancient positive selection in humans (but not HARs) were enriched in replication variant regions (focusing on variants in iPSC – the cell type better reflecting the germline; Figure S7 C, D, F). On the other hand, loss of function intolerant genes, as well as all protein coding genes, were found to be significantly depleted in iPSC replication timing variant regions (Figure S7 B, E). Taken together, these results suggest that replication timing alterations are unfavorable at conserved regions, possibly because they have an impact on genome function. Conversely, sequence divergence appears to be associated with replication timing differences between species.

### A complex association between DNA replication timing and gene regulation

Since replication timing is correlated with genome regulation (e.g. gene expression, chromatin accessibility; Figure 1), we tested whether replication timing variation was itself correlated with differences in gene expression or chromatin accessibility. Indeed, replication timing differences were positively correlated with gene expression variation (LCL: r=0.22, iPSC: r=0.25) and most replication timing variants (LCLs: 407/731, 56%, z-test p=7.0×10^-4^; iPSCs: 312/557, 56%, z-test p=9.2×10^-4^) contained predominantly genes with inter-species gene expression variation that corresponded to the direction of replication timing variation (i.e. earlier replication associated with elevated gene expression, later replication with reduced gene expression) (Figure 2H; S5 A). Similarly, we observed a positive correlation between replication timing variation and chromatin accessibility, assessed using ATAC-seq (LCL: r=0.35) and the histone modifications H3K27ac (LCL: r=0.59; iPSC: 0.53), H3K4me1 (LCL: r=0.44), and H3K4me3 (LCL: 0.51): the earlier replicating species had a relatively higher density of the open chromatin marks compared to the later replicating species (Figure 2 I-J; S4 F-G; S5 B). In contrast, density of the repressive chromatin mark H3K27me3 was not significantly correlated with replication timing variation (LCL: 0.05; iPSC: −0.09; Figure S4 H; S5 C). Overall, 90% of autosomal human-chimpanzee replication timing variant regions had concordant changes in replication timing and either gene expression or chromatin structure (based on H3K27ac, the histone mark most correlated to replication timing), and 52% (343/656) had concordant changes in all three.

To get a better understanding of the cause-and-effect relationships between DNA replication timing and chromatin, we analyzed their spatial co-variation. In some variant regions, chromatin structure differed between species primarily at the source site of replication timing variation (Figure 2F, top left; 2G, top right), suggesting that chromatin structure could be a determinant of the observed replication timing variation. In contrast, in other instances, differential chromatin structure/accessibility was present across the entire variant region (e.g. Figure 2G, top left and bottom left), suggesting that instead, replication timing could be exerting long range effects on chromatin structure. Similarly, there was no consistent spatial relationship between replication timing and gene expression variation; in some cases, gene expression varied concordantly primarily at the source site of replication timing variation (Figure 2F, bottom right; 2G, top right), while in other cases, concordant replication timing-gene expression variation extended across the entire variant (Figure 2F top right; 2G, bottom left). The evidence for each of these patterns across numerous genomic regions suggests that the interaction between replication timing and gene expression regulation is complex and locus-specific. As an extension, gene expression variation was not generally higher for genes in replication timing variant regions compared to non-variant regions (mean log FC of genes in variants=1.3, non-variants=1.4), together indicating that gene expression and replication timing variation, while often linked, are neither sufficient nor necessary drivers of one another.

### The genetic basis of replication timing evolution

The differences between species described above are suggestive of past and/or ongoing evolution of replication timing. As an extension of this observation, ongoing evolution is expected to manifest as inter-individual variation within a given species. Indeed, we have previously shown that replication timing varies among humans at hundreds of genomic locations [16, 18]. Consistently, in the current LCL sample set we identified 185 human and 195 chimpanzee genomic regions with significant variation among individuals (Methods; Figure 3). Of those, 73 regions varied among individuals in both species, significantly more than expected by chance (18 expected; z-test p=6.5×10^-40^) (Figure 3 C, F, G), while 112 regions were variable only among humans and 122 only among chimpanzees (Figure 3 D, E, G). More than half of the intra-species variants were also identified as inter-species variants (Figure 3 D-G), including for variants that were shared across species (40/73; 22 expected; z-test p=7.8×10^-6^) or those that were species-specific (63/112 human; 34 expected; z-test p= 4.8×10^-9^; 57/122 chimpanzee; 34 expected; z-test p=1.1×10^-6^). When directly testing human-chimpanzee variants for within species variation, 239 variants were also polymorphic in at least one of the species. Notably, 20 resolved human-evolved variants were also variable among humans, suggesting ongoing evolution of human replication timing in these regions. Taken together, we find significant evidence for replication timing polymorphism within both humans and chimpanzees, a substantial fraction of which appears to represent deep evolutionary processes that manifest as either conserved replication timing variation (Figure 3C) or concomitant intra- and inter-species variation (Figure 3 D-F).

**Figure 3.**
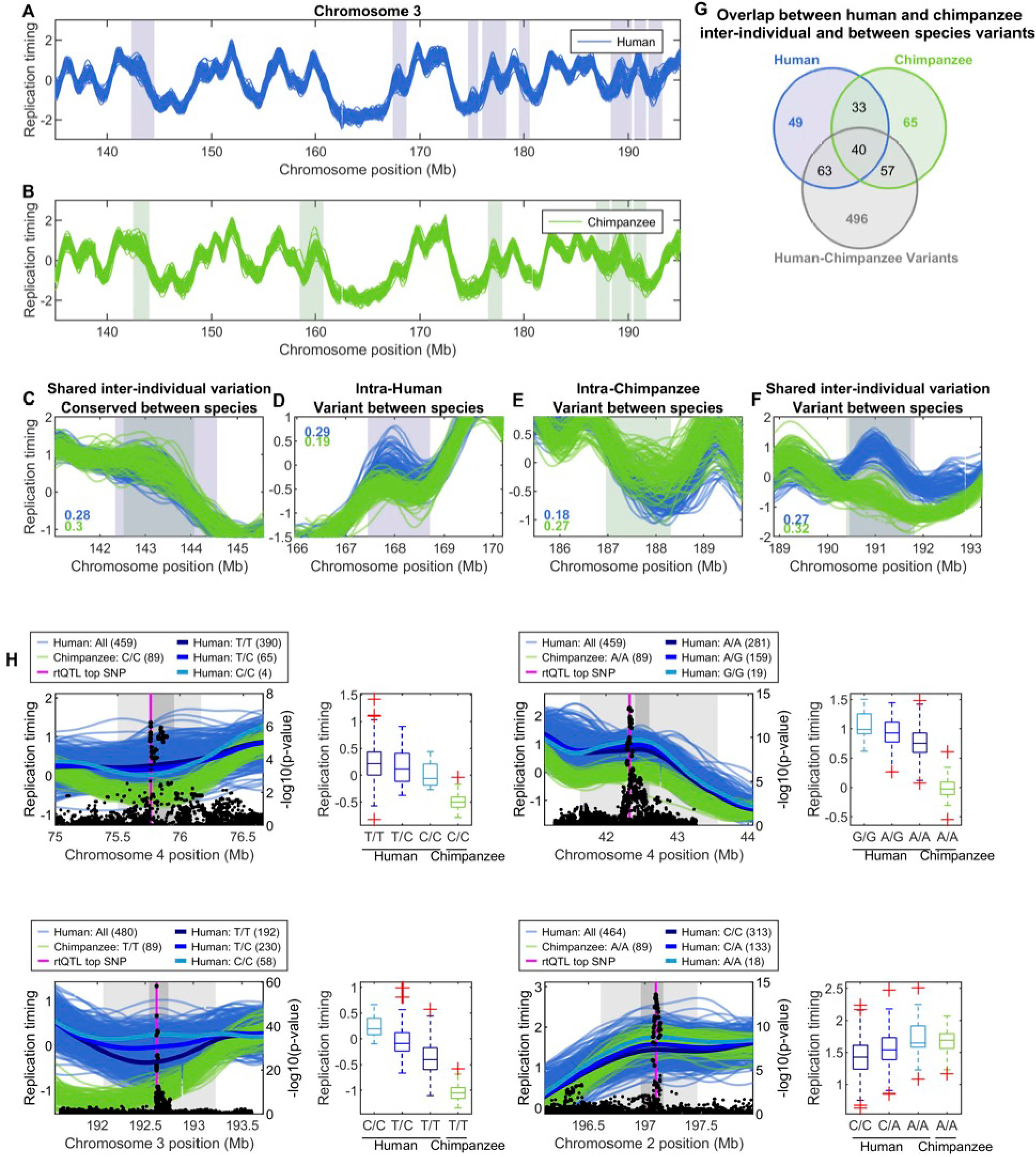
Genetic variation underlying inter-individual and inter-species replication timing variation. (A, B) Regions of inter-individual replication timing variation in human (A) and chimpanzee LCLs (B) for a section of chromosome 3. (C-F) Examples of shared and species-specific inter-individual replication timing variant regions. The numbers within each plot indicate the maximum standard deviation of replication timing values within the variant region for human and chimpanzee (blue and green, respectively). (G) Sharing of inter-individual replication timing variant regions between humans and chimpanzees and with between species (human-chimpanzee) variant regions. (H) Examples of human rtQTLs that overlap human-chimpanzee replication timing variant regions. The top rtQTL SNP (magenta) falls within, or near, the source site of replication timing variation. Human 1000 Genomes data was used for replication timing profiles and boxplots. In three of the examples, replication was earlier in humans, and the chimpanzee allele at the top associated rtQTL SNP matches the late replicating human allele. The opposite direction, i.e., derived late replication in humans, is observed in the bottom right example.

Identifying intra-specific replication timing variation is particularly relevant in the context of this study, since such variation can be used to map replication timing quantitative trait loci (rtQTLs; [16, 18]) which can then be tested for association with inter-species variation. Our population-level measurement of replication timing across species thus lends itself to the identification of the genetic basis of replication timing evolution.

To map rtQTLs in chimpanzees, we used fastQTL as recently described ([18]; see Methods), controlling for the population structure and relatedness of our sample set (Figure S3 C-F). This unbiased genome-wide analysis identified 21 rtQTLs – a relatively small number which we ascribe to the limited sample size and relatedness of the 89 chimpanzees. To increase rtQTL discovery power, we further mapped chimpanzee rtQTLs directly in regions of chimpanzee inter-individual replication timing variation and human-chimpanzee replication timing variation. This identified an additional 31 rtQTLs among the 195 chimpanzee inter-individual variants, and a further 33 in the 656 autosomal human-chimpanzee variant regions (Figure S8).

To compare replication timing polymorphisms to genetic variation in the current human LCL samples, we took advantage of the much larger number of 1,775 rtQTLs that we previously mapped in human stem cells [18] and 3,752 rtQTLs that we independently mapped in LCLs from the 1000 Genomes project (our unpublished results). We validated 276 of the 1000 Genomes rtQTLs directly in the current human samples (out of 2,793 rtQTLs for which we had all three genotypes at the top associated SNP location) and also verified that most human inter-individual variants from the current study overlapped rtQTLs in the 1000 Genomes dataset (141/185, Z-test p=8.3×10^-7^; 83 overlapped human rtQTLs validated in current samples). Thus, rtQTLs reflect the genetic basis of replication timing variation at a substantial fraction of the sites we mapped in this study.

Since we observed a high concordance of within species variation with between species variation (Figure 3 A-G), we predicted that human rtQTLs will also be associated with replication timing variation between humans and chimpanzees and could thus shed light on the genetic evolution of this form of variation. Indeed, 187 LCL and 57 iPSC human-chimpanzee variant source sites overlapped an rtQTL top associated SNP in the respective cell type, significantly more than expected based on randomizations (LCL: z-test p=3.5×10^-33^; iPSC z-test p=0.004).

Since some rtQTL associated SNPs affect replication timing at a distance, we also confirmed that source sites were enriched in rtQTL affected regions in addition to rtQTL SNPs per se (LCL: z-test p=7.5×10^-4^).

To test whether rtQTL sequences, at least in part, stand at the basis of replication timing evolution, we asked whether the derived allele matched the evolved replication timing state. For example, we would predict that humans carrying the ancestral allele for an rtQTL would have the ancestral replication timing (i.e. similar to chimpanzees), while humans with the derived allele would have the derived (i.e. different) replication timing state. We tested this prediction on human rtQTLs that spanned an inter-species variant source site and used the top associated rtQTL SNP and strongly linked SNPs (LD>0.8). We found a strong enrichment of rtQTLs where the human-derived allele was associated with the evolved replication timing state, while the ancestral allele was associated more closely with the chimpanzee replication timing (at least one tested SNP for 1,249/1,605 rtQTLs, 78%, permutations p=0.0072; >50% of SNPs for 741/1,605, 46%, p=0.0012; see examples in Figure 3H). Of these rtQTLs, 227 spanned human-chimpanzee variant regions that were resolved as changes in the human lineage. In 215 of these rtQTLs (95%), the chimpanzee allele of at least one tested SNP in high LD matched the macaque allele, suggesting that the genetic association may be sustained throughout the primate lineage as well.

The same analysis for the chimpanzee rtQTLs revealed that the chimpanzee derived allele matched the evolved replication timing state for 18 out of the 44 chimpanzee rtQTLs mapped in human-chimpanzee variants, suggesting that the derived chimpanzee allele was contributing to the difference in replication timing between humans and chimpanzees in these regions.

Importantly, 30 of the 44 chimpanzee rtQTL associated regions (mapped in human-chimpanzee replication timing variants) were also shared with a human rtQTL (expected 21, z-test p=0.005). This was not the result of ancient polymorphisms with conserved effects on DNA replication timing, as humans and chimpanzees did not share the associated rtQTL SNPs. Instead, this suggests that independent genetic contributions influence the replication timing of a given region across species, while rtQTL sharing further reflects either evolutionary pressures to maintain replication timing polymorphisms, or relaxed selective constraints to fix replication timing at these loci.

### Shared genetic causes of replication timing and gene expression evolution

We showed above that the evolution of DNA replication timing can be ascribed to sequence evolution while it also impacts regulatory evolution. Considered jointly, and further with the sequence determinants of gene regulation, these observations could potentially reveal how gene regulation and DNA replication timing have co-evolved. We previously showed that, across humans, replication timing and gene expression variation often share genetic causes [18]. We thus took advantage of comprehensive mapping of gene expression QTLs (eQTLs) in LCLs by the GTEx consortium [35], and compared them to the top associated SNPs (and/or SNPs in LD>0.8 to that top SNP) of the rtQTLs we found to be associated with replication timing variation between humans and chimpanzees. We found 488 rtQTLs (out of 1,605 that overlap human-chimpanzee variants) were also significant eQTLs (q-value<0.05; 192 unique variant regions). At these eQTLs, 64% of the involved genes (194 out of 301 for which expression data was available) had concordant changes in gene expression and replication timing, suggesting shared genetic causes of replication timing and gene expression evolution.

A notable example was a human-chimpanzee variant region that was both an rtQTL (Figure 4A) and an eQTL for two protein coding genes (Figure 4C) and one lincRNA. The rtQTL top SNP was the same as the top eQTL SNP (rs7806550), and there were no SNPs within 10 Kb of the variant with LD>0.4 (Ensembl 1000 Genomes YRI LD). Among human populations, the ancestral allele frequency for rs7806550 was highest in African populations (19%) and much lower in out of Africa populations (0-4%) (Figure 4B). This suggests that the derived allele emerged in the common ancestor of humans and increased in frequency to become the major allele in modern day humans. The region impacted by this shared rtQTL-eQTL was earlier replicating in humans than chimpanzees, and the two protein coding genes associated with the eQTL, ITGB8 (integrin complex subunit that mediates cellular interactions) and MACC1 (regulator of hepatocyte growth factor receptor involved in cell growth and motility), were also more highly expressed in humans (Figure 4A). Although rs7806550 is not known to be associated with any human phenotype (GWAS catalog; [36]), it fell within a strong LCL enhancer, and was predicted to affect two transcription factor binding motifs – for GATA and for HDAC2 – where the alternate allele (T; also ancestral allele) has higher binding affinity (Figure 4D). HDAC2 catalyzes deacetylation of lysine residues at the N-terminal regions of core histones (H2A, H2B, H3 and H4) and we previously showed that HDAC2 binding is associated with late replicating rtQTL alleles [18]. In this specific example, the ancestral allele (with higher HDAC2 binding affinity) was later replicating and matched chimpanzee replication timing. These observations can be explained if the human derived allele interrupts the HDAC2 binding site, decreases its ability to bind and deacetylate histones in the area, thus leading to greater histone acetylation, greater chromatin accessibility, and ultimately earlier replication and higher expression levels of genes in the immediate area. Interestingly, we also identified this region as variant between human and chimpanzee iPSCs (Figure S5 D, middle), and previously showed it to be the location of a human iPSC rtQTL [18]. Overall, this indicates that sequence changes may coordinate the concomitant evolution of replication timing and gene expression, through a chromatin intermediate.

**Figure 4.**
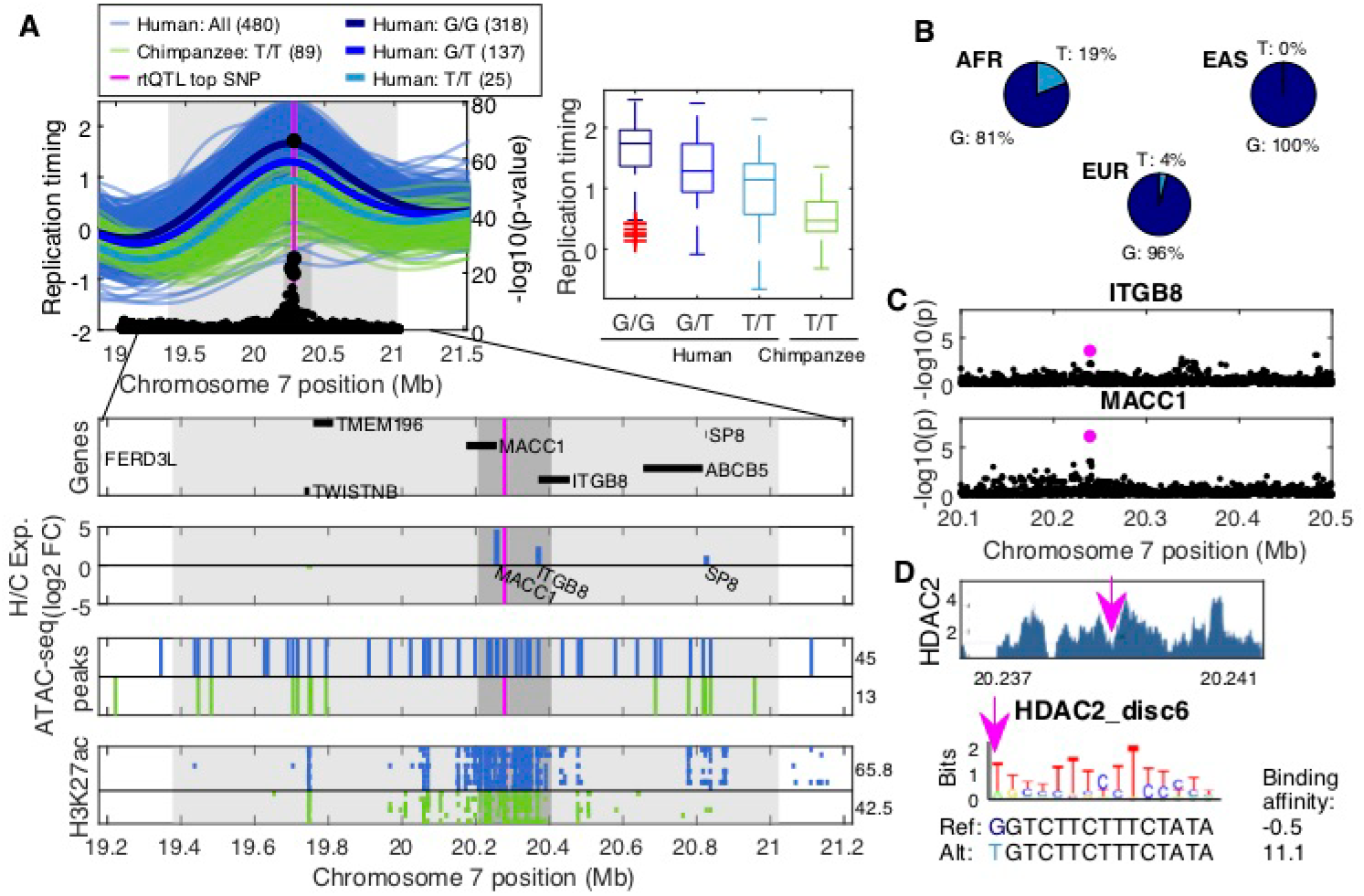
A genetic variant affecting HDAC2 binding, DNA replication timing and regional gene expression. (A) Chimpanzee replication timing profiles (this study) together with all replication timing profiles from the 1000 Genomes African super-population (AFR) and averaged replication profiles per genotype at the top associated rtQTL SNP (rs7806550). Genes, expression levels, ATAC-seq and H3K27ac density shown below. Magenta: rs7806550, located between the genes ITGB8 and MACC1. (B) AFR, East Asian (EAS), and European (EUR) 1000 Genomes phase 3 allele frequencies for rs7806550. (C) ITGB8 and MACC1 LCL eQTLs (GTEx). Magenta: rs7806550. (D) A HDAC2 regulatory motif is altered by the rtQTL-eQTL top SNP (rs7806550), with the alternate allele (T) having higher binding affinity. Top: Encode HDAC2 ChIP-seq data for GM12878 [37]. Bottom: Sequence logo of HDAC2 regulatory motif that is altered by rs7806550 (magenta arrow). Binding affinities are from HaploReg [38] (Δ affinity = 3,104-fold).

## Discussion

A long-standing question in human biology is what are the genetic changes that distinguish us from other species? The significant sequence similarity between humans and our ape relatives has pointed to regulatory evolution as a likely explanation of our unique phenotypes [3, 4]. However, studies of the evolution of gene expression and other epigenetic features has fallen short of fully explaining the complex adaptations in the human lineage. An understudied biological process from an evolutionary standpoint has been DNA replication timing, a fundamental genomic process that bridges genome regulation and maintenance. Previous studies of replication timing evolution have been restricted by small sample sizes, limiting their ability to fully describe evolutionary alterations and reveal the genetic drivers and the impacts of replication timing evolution. This has also limited the understanding of the forces that drive replication timing evolution and thus understanding of its functional significance. Here, we utilized population-scale replication timing profiling of humans, chimpanzees, and rhesus macaques to identify hundreds of genomic locations that vary in replication timing within and between these species, regulatory features that co-vary with replication timing across species, and sequence variants associated with replication timing evolution.

Notwithstanding local variation, the majority of the genome exhibits highly conserved replication timing both between and within species. This is unlikely even for lack of input sequence variation, as we have previously shown that multiple sequence determinants, spread over areas spanning several megabases, can influence the activity of any given replication origin [18]. For the replication timing variants that we do observe, most are quantitative (changes in replication origin firing times), and none can be considered of very large magnitude (e.g., >half of S phase). Thus, it appears that DNA replication timing is largely under evolutionary constraint and thus likely harbors an essential function(s), consistent with previous studies [13, 14, 39]. On the background of this overall conservation, we provide evidence for replication timing evolution across more than 30% of the human genome, including 123 genomic regions that have specifically evolved in the human lineage. This extent of inter-species replication timing differences is on par with gene expression evolution (e.g., 24% of genes in LCLs; [24]) and far exceeds the ~1% sequence divergence between humans and chimpanzees. This illuminates the functional potential of DNA replication timing in human evolution, with sequence alterations that affect replication timing carrying a disproportional effect on the genome compared to other regulatory sequence adaptations. The null assumption should be that these alterations are evolutionary neutral, which would be consistent with the observation that replication timing evolution mimics the phylogenetic tree of the same studied species. Shared variation between and within species, and shared rtQTLs (similar to previous observations of shared eQTLs; [40–42]), could also be a reflection of neutral drift and lack of local constraint, although an intriguing alternative possibility in this case is the action of balancing selection. Nonetheless, we did identify numerous genes with evidence of having undergone positive selection in humans as implicated in inter-species replication timing variant regions. This, and the general correlations with variation in gene regulation and potentially with mutation rates (see further below), points to the possibility that a subset of replication timing-evolved regions carry important functional implications for human evolution.

Studying a large number of individuals provided the unique ability to detect replication timing variation concomitantly between and within species. We observed extensive overlap of variation within and between species, pointing to deep and ongoing evolutionary processes impacting replication timing. This overlap also highlights the value of large sample sizes in evolutionary studies of replication timing, since variation between species is likely obscured in studies using smaller sample sizes. Furthermore, identifying and comparing variation within and between species enabled us, for the first time (to our knowledge), to reveal some of the genetic determinants of human replication timing evolution. We did this using an rtQTL mapping approach, which we applied to each species separately and then combined by considering co-occurring inter-species replication timing variation and linking derived rtQTL alleles to derived replication timing states. We anticipated that a much larger fraction of replication timing variants is determined by sequence evolution and could have been revealed with a larger samples size and hence greater power to detect rtQTLs.

Identification of genetic determinants of replication timing evolution provides a means for revealing mechanisms of replication timing control as well as the population genetic and evolutionary dynamics of replication timing. A notable example we highlighted pertains to the role of histone acetylation. HDAC binding has been described previously as a repressor of replication timing [18, 43], likely by promoting a more repressive chromatin state. Here, we showed that a sequence polymorphism impacting an HDAC2 binding site within a human LCL enhancer has likely led to variation in both replication timing and gene expression within humans as well as between humans and chimpanzees. The high frequency of the derived allele in humans suggests a likely directional evolution of this variant and therefore of replication timing in this case.

More generally, and consistent with previous studies, we observed correlated co-variation of DNA replication timing, chromatin accessibility and gene expression. A notable advantage of our study, however, is the ability to interrogate these relationships across hundreds of genomic regions harboring natural variation in DNA replication timing. While previous studies typically described these correlations as indicative of unidirectional causality relationships (in either direction; [12, 44–48]), our study suggests a more complex picture of replication timing and gene expression co-varying and potentially affecting each other (with chromatin structure being a likely intermediary) in a cell type-, locus- and context-specific manner.

Another important functional aspect of replication timing is its influence on the mutational landscape and therefore on local sequence evolution. Beyond validating the correlation between replication timing and rates of mutation and sequence variation, we found that sequence divergence between human and chimpanzee was elevated specifically in replication timing variant regions (in iPSC in particular). This suggests that evolutionary changes in replication timing could potentially alter local mutation rates and patterns. This would also be consistent with the observation that protein-coding genes (especially loss of function intolerant genes) were generally depleted in the same variant regions; thus, replication timing alterations in regions with highly conserved genomic function may be unfavorable, possibly due to the dual relationship of replication timing with genome regulation and genome maintenance.

More comprehensive mutational data would be required in order to test with sufficient statistical power how replication timing affects sequence evolution. The ability to obtain additional replication timing and mutation data for chimpanzees is, however, notably limited by the scarcity and regulatory limitations of using chimpanzee material. An alternative is to study replication timing variation within the more limited evolutionary timescale of human populations, or the broader timescale of more diverged mammalian species such as rodents compared to primates.

Another critical observation is that replication timing evolution is predominantly cell-type-specific. iPSCs and LCLs had very little overlap of inter-species replication timing variants. By inference, other cell types can be expected to show replication timing evolution at yet other genomic locations, and therefore the full functional impact of replication timing evolution would only be possible to evaluate in a larger number of cell types.

Overall, our findings highlight the importance of replication timing evolution as both a driver and consequence of sequence and regulatory evolution. DNA replication timing may thus carry an important yet previously under-considered role in human evolution. As such, it would be highly informative to incorporate replication timing in future studies of sequence and epigenetic evolution.

## Methods

### Sample preparation and whole genome sequencing

Genomic DNA from ninety chimpanzee lymphoblastoid cell lines (LCLs) (Table S3) was sourced from the Coriell Institute (Camden, NJ). These cell lines were originally derived at the Yerkes Primate Center in Atlanta, GA (samples from animals received prior to 2015). Genomic DNA from 85 human LCLs was sourced from the Coriell Institute and three LCLs from Grant Stewart (Table S3). Twenty-three rhesus macaque LCLs and a panel of seven human and seven chimpanzee induced pluripotent stem cell lines (iPSCs) were obtained from the Gilad Lab at the University of Chicago (Table S3) [22]. LCLs were cultured in Roswell Park Memorial Institute 1640 media with 15% fetal bovine serum, and iPSCs were cultured in mTeSR Plus media. Approximately one million cells of each cell lines were flash-frozen, and genomic DNA was extracted from them using the MasterPure DNA Purification kit (Epicentre, Madison, WI, USA). Genomic DNA from one additional human iPSC (AG25370) was obtained from the Coriell Institute.

All DNA samples were sequenced to approximately 20x coverage using the Illumina HiSeq X Ten with 2×150 paired-end reads (GENEWIZ, Inc., South Plainfield, NJ, USA). Chimpanzee LCLs were sequenced across two separate batches with two samples re-sequenced across batches as a control (NS03621 r=0.99; NS03639 r=0.97); clustering of replication timing values did not show evidence of a significant batch effect (batch 1 vs. 2 mean r=0.92). Human LCLs were sequenced in either batch primarily for other purposes, but used as batch-matched controls for this project. Rhesus macaque LCLs were sequenced in a third batch, with two chimpanzee LCL samples re-sequenced across batches as controls. All iPSC samples were sequenced in the same batch.

We were not able to generate reliable replication timing profiles for two chimpanzee LCLs, one chimpanzee iPSC, and two human iPSC samples. These samples had much lower mean correlation to the rest of the samples (chimp LCLs r=-0.03, −0.60; human iPSCs r=0.76, 0.83; chimpanzee iPSC r=0.8) and were removed from further analysis. One of the low-quality chimpanzee LCL samples was sequenced separately to approximately 40x coverage and yielded good quality data, thus we included these additional sequence data for this sample in our analyses; this led to a total of 89 chimpanzee LCL samples that were used for further analysis.

### Chimpanzee relatedness and population structure

We used a modified genome analysis toolkit (GATK; v4.1.4.0) best practices pipeline [49] to call SNPs and indels across the 89 chimpanzee LCL samples. We recalibrated base quality scores using chimpanzee dbSNP locations (prior to recalibration, we converted the dbSNP locations from the panTro5 to the panTro6 reference genome using the GATK tool LiftoverVcf (Picard; http://broadinstitute.github.io/picard/)). After recalibration, we genotyped each sample separately, then joint-genotyped all samples, as per the best practices. This resulted in a total of 28,476,465 variants including 23,911,740 SNPs and 4,125,969 indels. Next, we filtered the resulting variants with hard filtering thresholds based on recommendations from GATK (SNPs: QD<2.0, MQ<40.0, FS>60.0, SOR>3.0, MQRankSum<-12.5, ReadPosRankSum<-8.0; indels: QD<2.0, ReadPosRankSum<-20.0, InbreedingCoeff<-0.8, FS>200.0, SOR>10.0). This resulted in 26,492,303 total variants including 22,442,083 SNPs and 4,050,220 indels.

We used vcftools to evaluate the overlap of our called genetic variants with other datasets. Our variants overlapped 817,964/1,034,979 (79%) dbSNP variants, 9,375,559/25,923,958 (36%) variants from Prado-Martinez et al. 2013 [50], and 7,921,456/24,469,855 (32%) variants from de Manuel et al. 2016 [51].

Genotype PCAs were generated using the R package SNPRelate [52]. We pruned genotypes for minor allele frequency (<0.05), missing rate (>0.1) and linkage disequilibrium (>0.1) using snpgdsLDpruning before generating a PCA with snpgdsPCA.

We used KING [53] to calculate pairwise kinship values and infer relationships between chimpanzee LCL samples. Kinship values indicated a large number of first-degree relatives (77 pairs), thus we directly tested for the presence of first-degree familial relationships within our sample set. We first identified chimpanzee trios where two individuals had a first-degree relationship with the same third individual. A trio could represent several configurations such as two parents and their shared offspring, a three-generation family, or one parent and their two offspring. We isolated 18 two-parent and shared offspring configurations by assessing Mendelian error rates among high population frequency SNPs: two parents homozygous for the same SNP should not produce a heterozygous offspring; however, this pattern can occur for the three-generation and parent with two offspring configurations.

### Chimpanzee de novo mutation calling

We utilized the chimpanzee trios to call de novo (germline, or somatic cell line) mutations in the offspring of the trios and evaluate their relationship with replication timing. Genotyping, mutation identification, and candidate mutation filtering followed a pipeline we describe in detail elsewhere (Caballero et al., in preparation), modified for chimpanzee genomic resources. Briefly, BAM files were recalibrated with GATK using chimpanzee dbSNP. We did not recalibrate genotypes due to the lack of training resources. Candidate mutations were removed around the HLA locus (chr6:28,000,000–33,600,000 in panTro6). To remove inherited variants where the genotype was miscalled in a parent, we removed candidate mutations where any other LCL chimpanzee sample in this study contained reads matching the mutant allele. After all filtering steps, 14,774 autosomal mutations remained in the 18 LCL offspring (mean: 820.77, range: 273-1,439). We expect the majority of these to be cell line mutations, and a small number to be germline mutations [9].

### Generation of DNA replication timing profiles

Human, chimpanzee and rhesus macaque whole genome sequencing data were aligned to their respective reference genomes (hg19, panTro6 and rheMac10) using BWA-MEM. We calculated GC-corrected sequencing read depth in 1 Kb uniquely alignable windows across each species’ genome, as previously described [17]. We merged 1 Kb to 10 Kb windows and then filtered the data as follows: for iPSC samples, we filtered out windows with CNVs and outlier data points using segmentation (MATLAB function *segment*) as previously described [17]. For the LCL samples, instead of segmentation we used a “population” filtering method similar to the one we described previously [18]. We calculated the median value (across samples) of each genomic window. We then used the median of these values across windows to represent the “common” copy number of the genome. Any window with a median of more than 0.4 copies (0.2 copies for male X and Y chromosomes) above or below this common number was removed. We repeated this “population” filtering method using the 25% percentile and separately the 75% percentile instead of median, which allowed to better capture outliers.

We further removed, in individual cell lines, genomic windows that were copy number outliers in specific samples (rather than across all samples). We removed data points that were at least 0.6 copies (0.4 copies for male X and Y chromosomes) above or below the common copy number (see above), or at least 0.35 copies (0.25 copies for male X and Y chromosomes) above or below the median copy number across samples of that specific replication timing window, in any particular sample. Together, these two parameters ensured the efficient filtering of absolute or relative outliers, respectively. For all samples (LCLs and iPSCs), we removed large (mostly chromosome level) copy number alterations, short (<500 Kb) segments between genome gaps and short (<100 Kb) segments between runs of missing data.

The filtered data was then smoothed between gaps ≥50 Kb and regions separated by ≥100 Kb with a cubic smoothing spline (MATLAB function csaps; parameter=10^-17^) and subsequently normalized to a mean of zero and standard deviation of one.

We generated consensus replication timing profiles for each species per cell type by averaging the filtered data across samples (before smoothing). We then smoothed and normalized the averaged filtered data with the same parameters as described per sample (see above).

### PC-correction of LCL profiles

We performed principal component analysis of raw filtered replication timing data for human, chimpanzee, and rhesus macaque LCLs separately and corrected each species’ data for 10 principal components (PC10) using linear regression [18]. The X chromosome for each species was corrected for male and female samples separately. We did not PC-correct the female macaque X chromosome since there were only three samples. We then smoothed and normalized the data as described above. We used the PC10 smoothed data for all analyses. We did not PC-correct the iPSC samples due to their low number.

### Generation of G1/S replication timing profiles

G1/S replication timing profiles were generated as previously described [9]. Briefly, approximately one million G1 and S phase cells for macaque LCL sample 76-06 and chimpanzee iPSC sample C3649 were sorted using a FACSAria Fusion (BD Biosciences, San Jose, CA, USA), DNA was extracted and sequenced as above. Following sequence alignment, we defined genomic windows of varying size, each encompassing 200 reads in the G1 data. We then counted the number of S phase reads that fell into those windows. Data was normalized to a mean of zero and standard deviation of 1. Windows of 100 Kb with standard deviation greater than 1.1 were removed as well as data points greater or less than 3.5 standard deviations. Filtered data was smoothed with a cubic smoothing spline (MATLAB function *csaps;* parameter=10^-17^).

### Comparison of WGS with Repli-seq

Chimpanzee lymphocyte and H2 Human iPSC Repli-seq data was downloaded from Replication Domain (https://www2.replicationdomain.com/database.php; Accessions: Int10455570, Ext30484475). The chimpanzee data was smoothed (MATLAB function *csaps;* parameter=10^-17^) and normalized to a mean of 0 and standard deviation of 1. We then used the UCSC liftOver tool to convert genomic coordinates from hg38 to panTro6 (hg38.panTro6.rbest.chain) for comparison with our data. During this conversion, 6,533/419,622 (1.6%) of windows were lost. Linear interpolation was used to match genomic window coordinates prior to calculating the correlations with the chimpanzee LCLs in this study.

### Replication timing window lift over

To compare replication timing profiles between species, we used the UCSC genome browser liftOver tool with the reciprocal best mapping chains to convert the center coordinate of the 10 Kb chimpanzee (panTro6) and rhesus macaque (rheMac10) replication timing windows to human (hg19) coordinates (panTro6.hg19.rbest.chain and rheMac10.hg19.rbest.chain, respectively). During this conversion, 6,615/270,666 (2.4%) of chimpanzee replication timing windows and 37,713/274,457 (13.7%) of rhesus macaque windows were lost.

### Replication origin prediction

We predicted the most likely locations of replication origins based on the sharing among samples of peaks in the replication timing profiles. We called local maxima in each sample, then used hierarchical clustering with average linkage and a distance threshold of 300 Kb to identify clusters of recurrent nearby peaks across all samples of a given cell type (e.g. human, chimpanzee, rhesus macaque LCLs). We removed replication origins that were present in less than 10% of samples in at least one of the species. We further removed origin calls that occurred within 100 Kb of a mapped structural variant (SV) [2, 21] or a gap in the human genome.

### Replication timing variation between humans and chimpanzees

To identify genomic regions with significant replication timing variation between species, we performed ANOVA tests comparing all samples from one species with all samples from the other species, in 200 Kb windows, sliding by 50 Kb, across all autosomes and the male and female X chromosome separately. Tested windows were considered to be significant if they passed a Bonferroni-corrected p-value threshold of 8.7×10^-7^. Overlapping significant windows were then merged, and p-values were recalculated. These were considered as “initial variant regions”. We excluded individual replication timing windows (10 Kb) from within these initial variant regions if they spanned genome gaps or that had a mean difference in replication timing between species of less than 0.2 standard deviations. These filters resulted in some of the initial variant regions being split or removed completely, yielding “filtered variant regions”. Filtered variant regions that were less than 200 Kb long were removed. In cases in which adjacent filtered variant regions had intervening replication timing windows with a mean replication timing difference between species greater than 0.2 (even if they were not significant in the initial ANOVA scan), we extended and merged these filtered variant regions, then recalculated the p-values of the merged variants. These were considered as “extended variant windows” and used for downstream analyses.

Since there is only one copy of the male X chromosome, we divided the filtering and extension thresholds above by 2 (i.e. used 0.1 standard deviation).

To classify the likely molecular type of replication timing variants, we tested each variant for overlap with predicted replication origin locations (peaks in the replication profiles) in each species. Variants harboring peaks (in >25% of samples) in both tested species were considered to be alterations in replication origin activation time, while variants with a peak in only one species (in >25% of samples) were considered to be an evolutionary gain or loss of a replication origin (more details below).

Next, we utilized the predicted origin locations to identify the most likely “source” sites of replication timing variation within each variant. We identified the called origins within each replication timing variant and considered only those that were present in >25% of samples of the species with earlier replication (including shared origins, in which both species had an origin in >25% of samples). Most variant regions contained only one origin, which was then considered as the source site. In cases with more than one origin within a variant region, if there was a valley in either the human or chimpanzee consensus profiles (or both, in which case the two valley locations were averaged) between the two origins, or otherwise if the origins were separated by more than 500 Kb, we split the variant region at the valley (or middle location, respectively) between the two origins and considered each origin to be a source site for its own variant. Otherwise, we considered the source site to be the middle location between the origins. In either case, the source sites were regarded as 200 Kb regions centered at the origin locations, but were not allowed to extend beyond the bounds of the variant region.

### Inference of directionality of evolutionary changes

To identify the specific lineage (human or chimpanzee) in which replication timing has likely evolved at replication timing variants, we compared each variant to the replication timing in rhesus macaques. This was done only for LCLs, for which we had data for all three species. For each variant, we calculated the pairwise Euclidean distance of consensus replication timing values between each species pair (i.e. human to chimpanzee, chimpanzee to macaque, human to macaque) at the source site and identified the species pair with the smallest distance. Variants for which the smallest distance was between humans and macaques were preliminarily considered to be evolutionary changes that occurred in chimpanzees, while replication timing variants that had the smallest distance between chimpanzees and macaques were preliminarily called as changes in humans. Replication timing variants for which the smallest distance was between humans and chimpanzees were considered unresolved in the absence of additional outgroups.

We then subjected the preliminary human and chimpanzee resolved changes to two quality filters. First, we required that the species pair with the smallest distance (see above) had replication timing profiles of similar shape. We assessed this for each preliminary resolved variant by calculating the correlation of consensus replication timing values within a 500 Kb window centered at the source site for the species pair with the smallest distance. If the correlation was less than 0.1, the region was re-categorized as unresolved. Second, we filtered the preliminary resolved changes where the species pair with the smallest distance (see above) was high (>1, or >1.5 for regions where the human to chimpanzee distance was greater than 3) or where the macaque consensus profile was equally distant to the human and chimpanzee consensus profiles (the difference between the human to macaque and chimpanzee to macaque distances was <0.35).

We manually filtered an additional 48 regions that had a visually different replication timing profile shape in macaque (despite passing the correlation threshold), or where the macaque profile visually looked equidistant from human/chimpanzee as they were from each other.

Preliminary resolved variants that passed the filters comprised the final resolved variants, which we then categorized into advances and delays. If the species in which the evolutionary change was inferred to have occurred was earlier replicating that the other two, the variant was called as an advance in that species, while delays were considered to be cases in which the species that underwent replication timing evolution was later replicating than the others. We then used the previously classified molecular type of replication timing variants (see above) to subset the advances and delays into gains and losses of origins versus changes in origin activation time. In addition, gains required that there was no origin in greater than 25% of macaque samples, while losses required an origin in more than 25% of macaque samples. Hierarchically-clustered heat maps of replication timing values across samples were generated with the MATLAB function *clustergram*.

### Replication timing variants shared between LCLs and iPSCs

To identify replication timing variants shared across cell types, we analyzed, for each human-chimpanzee LCL replication timing variant, the replication timing in human and chimpanzee iPSCs. We calculated the pairwise Euclidean distances of human and chimpanzee LCL and iPSC consensus replication timing values at each replication timing variant source site. Variants were considered to be shared across the two cell types if the distances within species (e.g. human LCL to human iPSC; chimpanzee LCL to chimpanzee iPSC) were lower than the distances within cell types (e.g. human LCL to chimpanzee LCL; human iPSC to chimpanzee iPSC). Of the shared variants, we also asked if the shape of the profiles was consistent within species by calculating the Pearson correlation of consensus replication timing values within species (e.g. human LCL to human iPSC; chimpanzee LCL to chimpanzee iPSC). Shared variants were considered to be of similar shape if the within-species correlations were greater than 0.5.

### Association with gene expression, chromatin accessibility, and sequence variation

Association between DNA replication timing and gene density, gene expression, chromatin accessibility (ATAC-seq and H3K27ac ChIP-seq), and sequence variation (SNP density, human-chimpanzee divergence, de novo mutations) was performed for human and chimpanzee LCLs and, separately, iPSCs when data was available. Data was obtained and prepared for analysis as follows:

We used the center coordinates (hg19) of Ensembl protein-coding genes. Gene expression analyses were based on published data for LCLs and iPSCs [21–24]. Chromatin data was obtained from the following sources: LCL ATAC-seq peak data [25], LCL histone modification ChIP-seq peak data [23], and iPSC ChIP-seq peak data [22]. All chimpanzee ChIP-seq data was originally in the panTro3 reference genome while ATAC-seq data was in panTro5; both were lifted-over to hg19.

Human and chimpanzee SNPs were obtained from dbSNP and filtered for coding sites. Divergent sites between human and chimpanzee were inferred from the Ensembl Enredo-Pecan-Ortheus (EPO) 12 primate multiple alignments (release 104) (n=32,301,278). We used liftOver to convert these sites from hg38 to hg19 (n=32,253,773 lifted). Replication timing windows with zero divergent sites were not considered in the following analyses.

For each of the genomic features described above, the following genome-wide analysis was performed: we binned replication timing windows into 30 equally portioned bins and counted the number of genomic features within each bin. Ten bins were used for de novo mutations, due to their small number. We subsequently compared the number of genomic features in each bin to the average replication timing value for that bin.

The following replication timing variant analysis was performed on each of the following data types (ATAC-seq, ChIP-seq H3K27ac, ChIP-seq H3K27me3, ChIP-seq H3K4me1, and ChIP-seq H3K4me3) separately, but will collectively be referred to as chromatin accessibility peaks. Prior to this analysis, we mapped human accessibility peaks back to panTro3 (ChIP-seq) or panTro5 (ATAC-seq) and removed human peaks that were not successfully lifted-over. We counted the number of peaks within each LCL or iPSC variant region and calculated log2 fold change in peak density normalized by the total number of human and chimpanzee peaks [log2((# human peaks in variant region/total # human peaks) / (# chimp peaks in variant region/total # chimp peaks))]. For comparison, we also calculated log2 fold change in peak density for 200 Kb windows across the genome. Log2 fold change in peak density was compared to the change in replication timing of the windows.

### Gene ontology, constraint, and selection

Gene ontology enrichment was performed using the PANTHER Overrepresentation Test (Released 2022-02-02) [54–56], separately on protein-coding genes that fell into LCL human-chimpanzee variants, iPSC variants, and protein-coding genes that were shared across the LCL-iPSC variants. Gene enrichment was calculated with the Fisher’s Exact Test and evaluated with 5% FDR.

We analyzed LCL and iPSC replication timing at 481 ultra-conserved elements (UCEs) across humans, mice, and rats [34], loss of function intolerant genes (gnomAD [33]), 2,701 non-coding human accelerated regions [31], human-chimpanzee divergent sites, all protein-coding genes, and regions under ancient positive selection in humans [32]. For each of these genomic features, we binned replication timing windows into 10 equally portioned bins and counted the number of genomic features within each bin. We then compared the number of genomic features in each bin to the average replication timing value for the bin. To evaluate if each genomic feature was enriched or depleted in replication timing variants, we randomized the locations of LCL and iPSC autosomal variant regions 100 times (size and replication timing matched, +/− 0.25 standard deviation) and calculated the total number of genomic features that each random region overlaps. The distribution of the total number of overlaps for each iteration was compared to the observed number of overlaps for each feature with a Z-test.

### Inter-individual replication timing variation

We identified genomic regions with significant replication timing variation among individuals of a given species as regions with a relatively high standard deviation (SD) of the replication timing data across individuals. We calculated SD across samples for each replication timing window across the genome (using replication timing data for each species own reference genome). To identify regions with the greatest regional SD, we first smoothed the SD values (MATLAB function csaps; parameter=10^-14^) and then called peaks in the smoothed SD profiles. We removed peaks with SD lower than the mean of all autosomal SD peaks. The remaining SD peaks were considered to be centers of inter-individual variant regions. We then extended these regions until the closest local SD minima (identified from the smoothed SD profiles similar to peaks) or until the SD equaled the genome-wide mean SD. We filtered any extended variant regions that spanned gaps or that were shorter than 200 Kb and performed a pairwise t-test on the remaining variant regions. For each tested variant region, we identified significant sample pairs using a Bonferroni corrected p-value threshold. We removed variant regions that resulted from a single sample causing the observed variation. Chimpanzee variants were lifted-over from panTro6 to human hg19 coordinates; 10/195 regions failed to be fully lifted-over. We considered the intra-species variants as shared between humans and chimpanzees if a variant from one species overlapped the center of a variant from the other species or vice versa. We additionally tested human-chimpanzee replication timing variants for intra-species variants directly by the pairwise t-test and following filtering steps as described above.

### rtQTL mapping and validation

Chimpanzee rtQTLs were mapped genome-wide, in human-chimpanzee replication timing variants, and within chimpanzee replication timing variants using fastQTL as in [18]. Briefly, smoothed replication timing data was used with 10 phenotype principal components (PCs) and 3 genotype PCs as covariates. In the genome-wide analysis, phenotype windows were chosen with a FDR<0.1, while phenotype windows for human-chimpanzee replication timing variants and within chimpanzee variants were chosen as the center variant window coordinate. For all three analyses, a SNP was identified as significantly associated with the phenotype if it belonged to a group of at least three consecutive SNPs with FDR<0.1. Merging and filtering of rtQTLs was performed as in [18].

1000 Genomes rtQTLs were mapped in six populations separately as a part of a separate study, using fastQTL as in [18].

Validation of 1000 Genomes rtQTLs in the human samples from this study was performed as in [18]. Briefly, we calculated Pearson correlation between the top associated rtQTL SNP and the replication timing value at the location with strongest association with the rtQTL. We only tested rtQTLs where we had all three genotypes of the top associated SNP in the human samples from this study (2,793/3,752 rtQTLs). rtQTLs were classified as validated if the p-value was less than 0.05 and had the same direction of effect.

We obtained the chimpanzee and rhesus macaque allele at each rtQTL top associated SNP and SNPs in LD>0.8 to the top SNP from the Ensembl EPO primate alignments. We performed the following allele directionality analysis only on human rtQTLs where the associated region spanned a source site of human-chimpanzee replication timing variation. For each rtQTL, we evaluated the allele effect (i.e. whether the human derived or chimpanzee allele was earlier replicating) for the top associated rtQTL SNP and SNPs in LD>0.8 to that top SNP. We then counted the number of rtQTLs where the allele effect was consistent with the direction of replication timing difference between species for at least one of the evaluated SNPs. For example, if chimpanzees were earlier replicating we asked if the chimpanzee allele matched the early replicating human allele, while if humans were earlier replicating we asked if the chimpanzee allele matched the late replicating human allele. To assess significance, we permuted the number of rtQTLs that we tested for allele direction. For each rtQTL tested, we compared the direction of replication timing difference between species to permuted rtQTL haplotypes (top associated SNP and SNPs in LD>0.8), then asked if the permuted SNP allele direction was consistent with the change in replication timing of the tested rtQTL. We counted the number of rtQTLs with at least one tested SNP with a consistent allele direction. We repeated these steps for 1000 permutations and used a z-test to calculate p-value.

The allele directionality analysis was also repeated on chimpanzee rtQTLs that were called in human-chimpanzee replication timing variant regions. Allele effect (i.e. whether the chimpanzee derived or human allele was earlier replicating) was only evaluated at the top chimpanzee rtQTL SNP.

Chimpanzee rtQTLs were classified as shared with human rtQTLs if the top associated human rtQTL location fell within the chimpanzee rtQTL associated region. Significance was evaluated with 100 randomizations and a z-test.

### Identifying shared eQTLs

To identify shared genetic causes of replication timing and expression variation, we located GTEx LCL eQTLs that shared a top associated rtQTL SNP (and/or one in LD>0.8 to the top associated SNP). Significant GTEx eQTLs had a q-value less than 0.05. For the shared rtQTL-eQTL example given, we used HaploReg [38] to identify if rs7806550 interrupted any regulatory motifs. Sequence logo and binding affinities for HDAC2 were obtained from HaploReg [38] and ChIP-seq data for HDAC2 was obtained from ENCODE [37].

## Acknowledgements

We thank Ana Rita Rebelo, Bronte Zhang, Lauren Mei, Sean Kim and Tiffany Ge for technical assistance, and members of our lab for critical reading of the manuscript. We thank Yoav Gilad for sharing iPS and LCL samples and for fruitful discussions. Chimpanzee LCL samples were sourced from the Yerkes Center (Grant No. P51OD011132); DONSON human LCL cell lines were a gift from Grant Stewart. This work was supported by the National Science Foundation (MCB-1921341 to A.K.). A.N.B. was partially supported by the Cornell Center for Vertebrate Genomics. A.D. is a Hunter R. Rawlings III Cornell Presidential Research Scholar.

## Data availability

Raw chimpanzee and rhesus macaque whole-genome sequencing data is available under SRA accession PRJNA856315, and human data is available under dbGaP accession phs002597. Three human LCL and six human iPSC samples were not consented for release of raw genomic sequence data. Processed replication timing data is available for all samples (Supplemental files 1 and 2).

## Notes

### Competing Interest Statement

The authors have declared no competing interest.

