## Supplementary Information for "The evolution of the human DNA replication timing program"

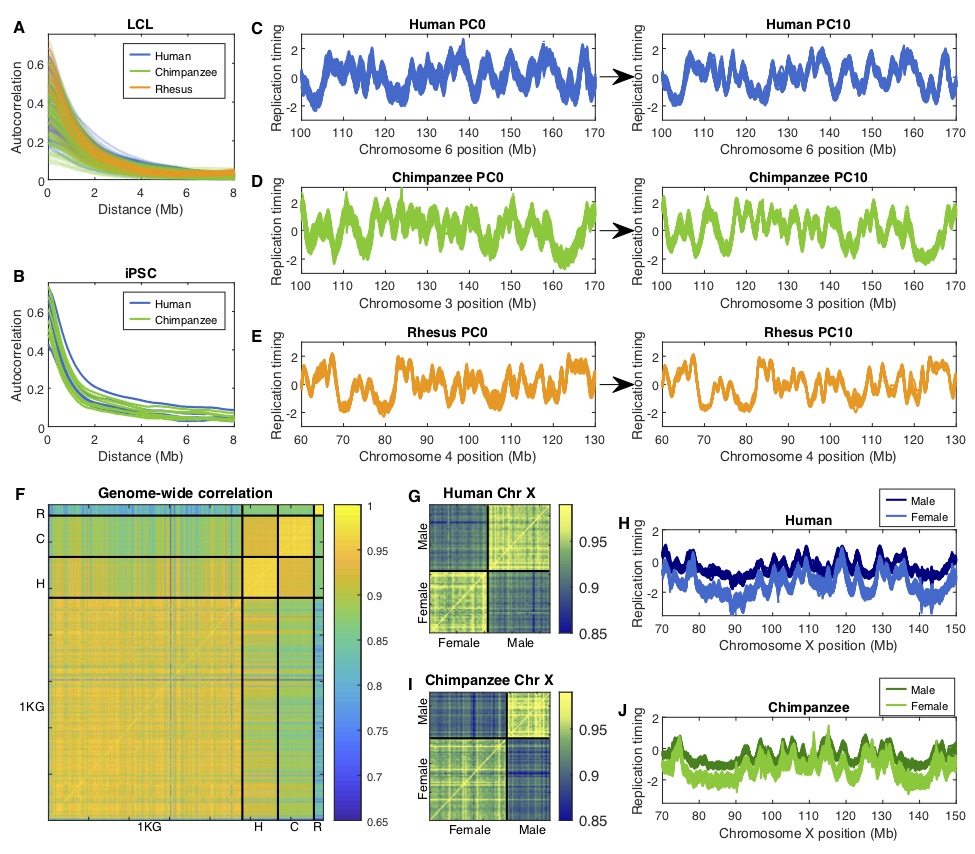

Figure S1. Quality control and data processing.

(A-B) Autocorrelation of raw autosomal replication timing data for LCLs (A) and iPSCs (B). (C-E) Smoothed replication timing profiles before principal component (PC)-correction (PC0; left) and after correction for 10 PCs (PC10; right) for human, chimpanzee and rhesus macaque LCLs. (F) Pairwise Pearson correlation of autosomal replication timing across all 1000 Genomes (1KG) African samples (n=480) to the human (H), chimpanzee (C), and rhesus macaque (R) samples from this study. Correlations are higher between 1000 Genomes samples and humans compared to chimpanzees or rhesus macaques from this study. (G, I) Pairwise Pearson correlation of human and chimpanzee LCL X chromosome replication timing values, confirming the conservation of sex differences. (H, J) Human and chimpanzee LCL X chromosome replication timing profiles for males and females. Female X chromosome profiles are later replicating than male, and also appear noisier (more diffuse).

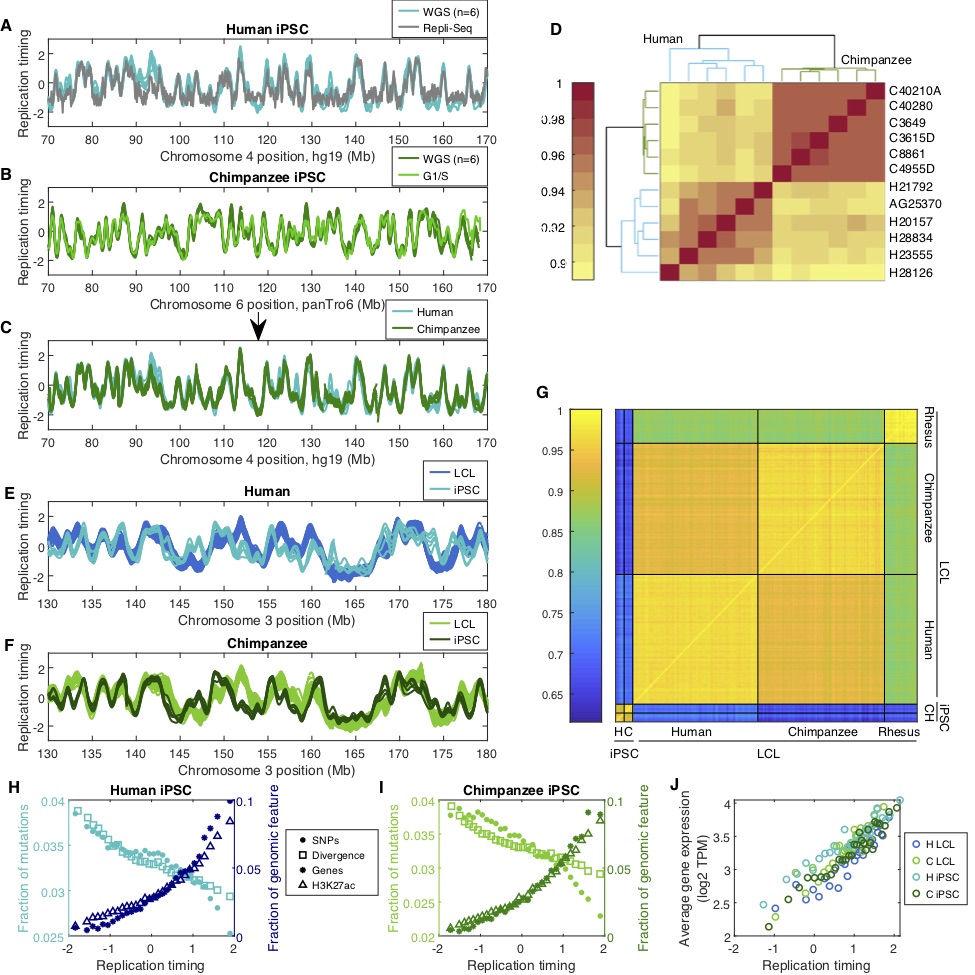

Figure S2. Replication timing varies between cell types more than between species.

(A-D, H, I) As in Figure 1, for human and chimpanzee iPSC data. Chimpanzee G1/S was generated in this study; human Repli-seq was obtained from Replication Domain (Accession: Ext30484475). (E-F) Overlaid comparison of human (E) and chimpanzee (F) LCL and iPSC replication timing profiles. (G) Pairwise Pearson correlation of autosomal replication timing across all iPSC and LCL samples. (J) Gene expression (averaged across cell lines for each cell type and species; data obtained from [1]) compared to replication timing for human and chimpanzee LCLs and iPSCs, at human-chimpanzee orthologous genes (with average TPM>0.1) in 30 replication timing bins.

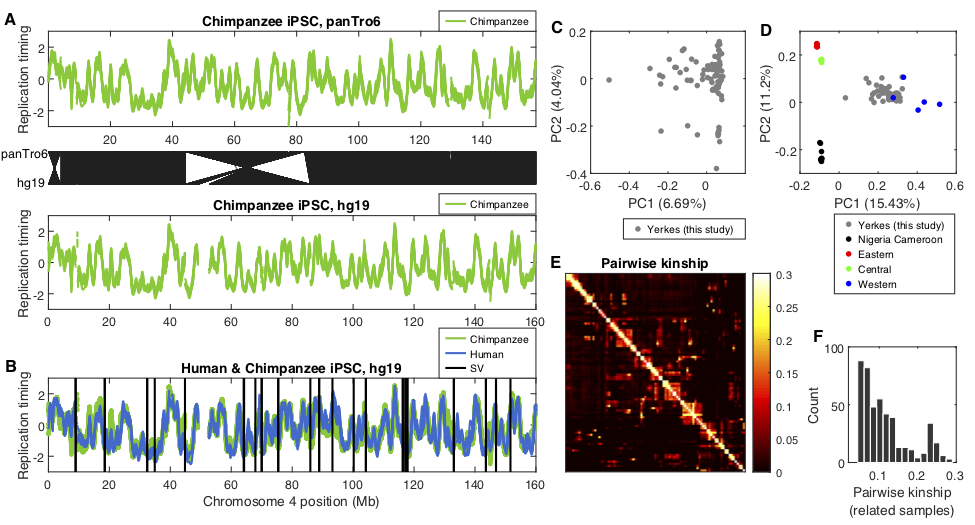

Figure S3. Chimpanzee relatedness, population structure, and replication timing coordinate conversion. (A) Chimpanzee iPSC chromosome 4 replication timing profiles in panTro6 (top) and human hg19 (bottom) coordinates. Black lines between the top and bottom panels indicate syntenic loci between panTro6 and hg19 genome builds. Two large inversions are apparent. (B) Human and chimpanzee iPSC replication timing profiles for chromosome 4 (hg19 coordinates) along with previously mapped structural variants (SVs; inversions, deletions, and insertions) [1, 2] larger than 100 Kb between human and chimpanzee. (C) Genotype principal component analysis (PCA) for the chimpanzee LCLs in this study. PC2 separates the samples into two groups (which does not correspond to sequencing batch). (D) Genotype PCA for chimpanzee LCLs in this study (“Yerkes LCLs”) together with samples from known chimpanzee sub-populations [3], indicates that the chimpanzee samples were primarily from the western chimpanzee sub-population (*Pan troglodytes versus*). (E) Pairwise kinship values across all samples. Values were clustered using hierarchical clustering. (F) Distribution of pairwise kinship for related samples (3^rd^ degree relationship or closer, kinship>0.04) (463 pairs).

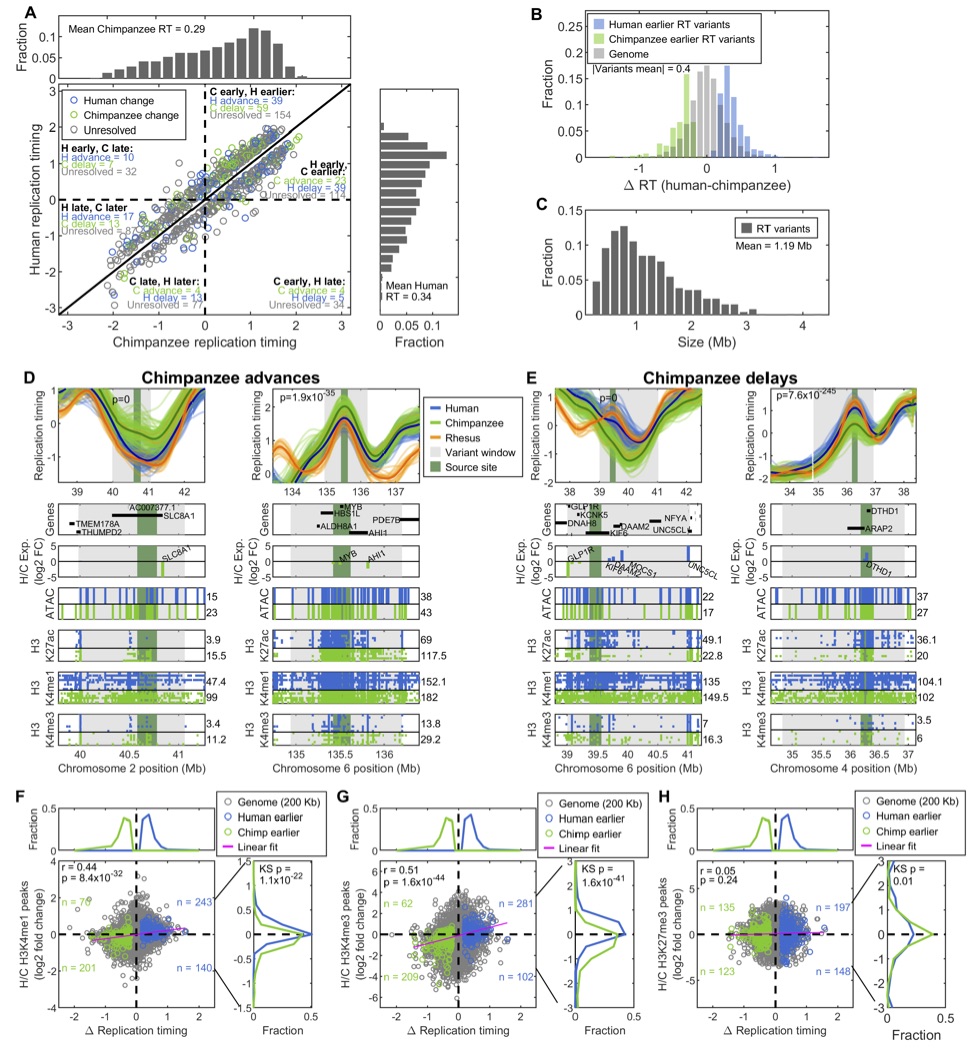

Figure S4. Further characterization of human-chimpanzee replication timing variants.

(A) Comparison of human and chimpanzee replication timing at variant regions (averaged data within each source site). Blue, green and gray data points represent variants resolved as having evolved in the human or chimpanzee lineages, or being unresolved, respectively. Top and right histograms: distributions of chimpanzee and human replication timing, respectively, within replication timing variant regions. (B) The distribution of the magnitudes of human-chimpanzee replication timing differences (averaged data within source sites of each variant). Background genome replication timing differences were calculated for each replication timing window across the genome. (C) Size distribution of human-chimpanzee replication timing variant regions. (D, E) Similar to Figure 2 F-G, for chimpanzee replication timing advances and delays. (F-H) As in Figure 2 J, but using human and chimpanzee LCL H3K4me1 (F), H3K4me3 (G), and H3K27me3 (H) ChIP-seq data.

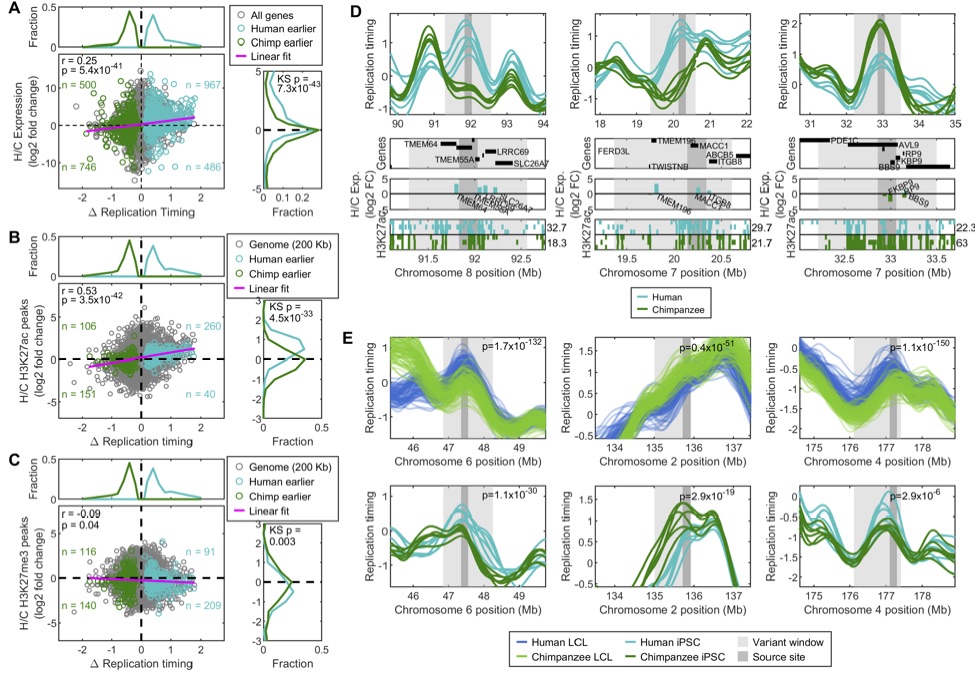
Figure S5. iPSC replication timing variants.

(A) Differences between human and chimpanzee iPSC replication timing compared to differences in gene expression, as in Figure 2H. (B-C) As in Figure 2J, but using human and chimpanzee iPSC H3K27ac (B) or H3K27me3 (C) ChIP-seq data. (D) Examples of iPSC human-chimpanzee replication timing variant regions. (E) Examples of human-chimpanzee LCL replication timing variant regions observed also in iPSCs. Top subplots show the LCL replication timing data and indicate the LCL variant window with p-value. Bottom subplots show the iPSC data for the same variant regions, p-values indicate the significance (ANOVA) of human-chimpanzee iPSC differences within the LCL variant region.

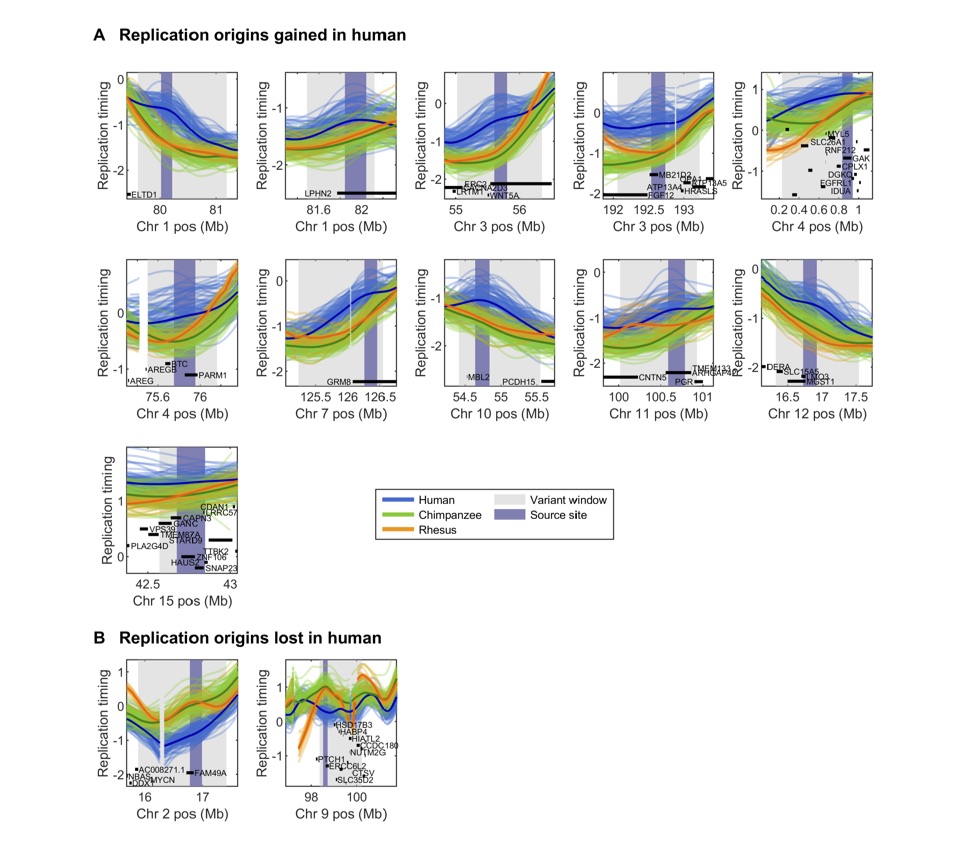

Figure S6. Gained and lost replication origins in humans.

Example regions with replication origins gained (A) and lost (B) in humans. Human-chimpanzee replication timing variant window and source site are indicated for each region along with genes that fall into each region. Some gene names removed for readability.

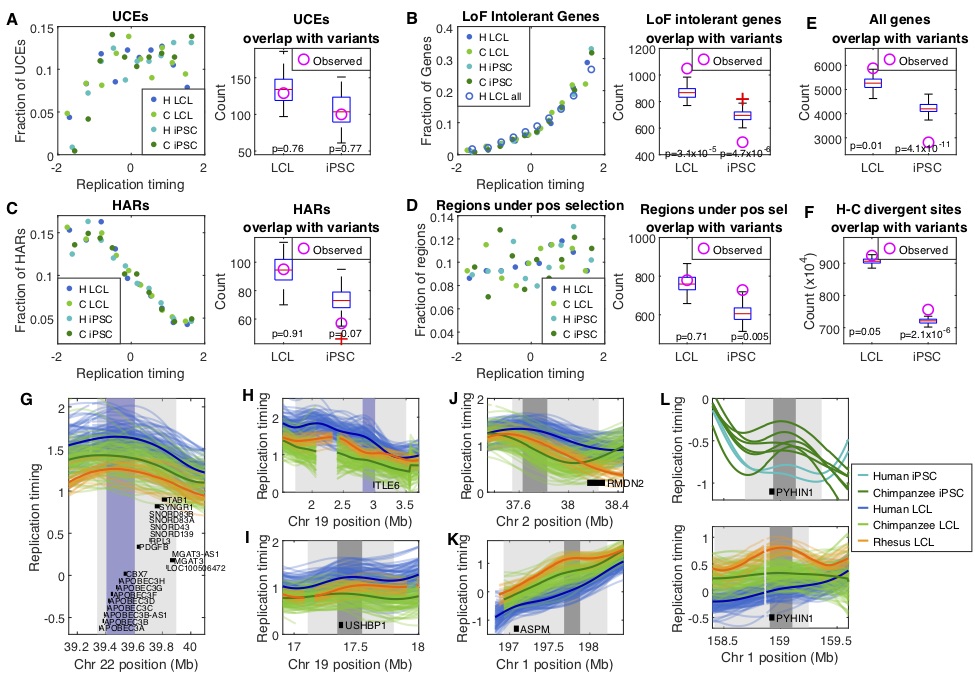

Figure S7. Replication timing at regions under constraint or adaptive evolution.

(A-D) Left subplots: fraction of ultra-conserved elements (UCEs, A), loss of function (LoF) intolerant genes (B), human accelerated regions (HARs, C), or regions under positive selection in human (D) in 10 replication timing bins compared to the mean LCL or iPSC replication timing of that bin per species. (B) Includes human LCL replication timing at all protein-coding genes for comparison to LoF intolerant genes. (A-D) Right subplots and (E-F): the overlap of each genomic feature with LCL and iPSC replication timing variants. Boxplots indicate overlap of the genomic feature with random genomic regions (number, size, and replication timing matched to the variants; 100 randomizations); magenta circle indicates the actual number of each genomic feature that overlaps the replication timing variants. P-values were calculated with a Z-test. (E-F) See Figure 1 for replication timing at all protein-coding genes (E) or human-chimpanzee divergent sites (F). (G) Human-chimpanzee LCL replication timing variant spanning the APOBEC gene cluster. (H-K) Examples of genes under adaptive evolution in humans that fall into human-chimpanzee LCL replication timing variants. (L) PYHIN1 falls within a shared LCL-iPSC human-chimpanzee replication timing variant.

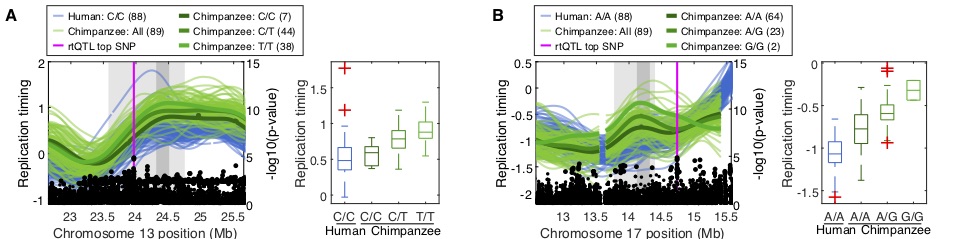

Figure S8. Chimpanzee rtQTLs.

(A-B) Examples of chimpanzee rtQTLs that overlap human-chimpanzee replication timing variant regions. Human data from this study was used for replication timing profiles and boxplots. In both examples, replication was later in humans, and the human allele at the top associated rtQTL SNP matches the late replicating chimpanzee allele.

Supplemental Tables and Files:

Table S1. Human-chimpanzee replication timing variant regions

Table S2. Gene ontology enrichment analysis

Table S3. Sample information

File S1. LCL replication timing data

File S2. iPSC replication timing data
